# Conformational changes in Apolipoprotein N-acyltransferase (Lnt)

**DOI:** 10.1101/497412

**Authors:** Benjamin Wiseman, Martin Högbom

## Abstract

In bacteria, lipoproteins are important components of the cell envelope and are responsible for many essential cellular functions. They are produced by the post-translational covalent attachment of lipids that occurs via a sequential 3-step process controlled by three essential integral membrane enzymes. The last step of this process, unique to Gram negative bacteria, is the N-acylation of the terminal cysteine by Apolipoprotein N-acyltransferase (Lnt) to form the final mature lipoprotein. Here we report 2 crystal forms of this enzyme. In one form the enzyme crystallized with two molecules in the asymmetric unit. In one of those molecules the thioester acyl-intermediate is observed. In the other molecule, the crystal packing suggests one potential mode of apolipoprotein docking to Lnt. In the second crystal form the enzyme crystallized with one molecule in the asymmetric unit in an apparent apo-state remarkably devoid of any bound molecules in the large open substrate entry portal. Taken together, these structures suggest that the movement of the essential W237 is triggered by substrate binding and could help direct and stabilize the interaction between Lnt and the incoming substrate apolipoprotein.

**Graphical Abstract:** 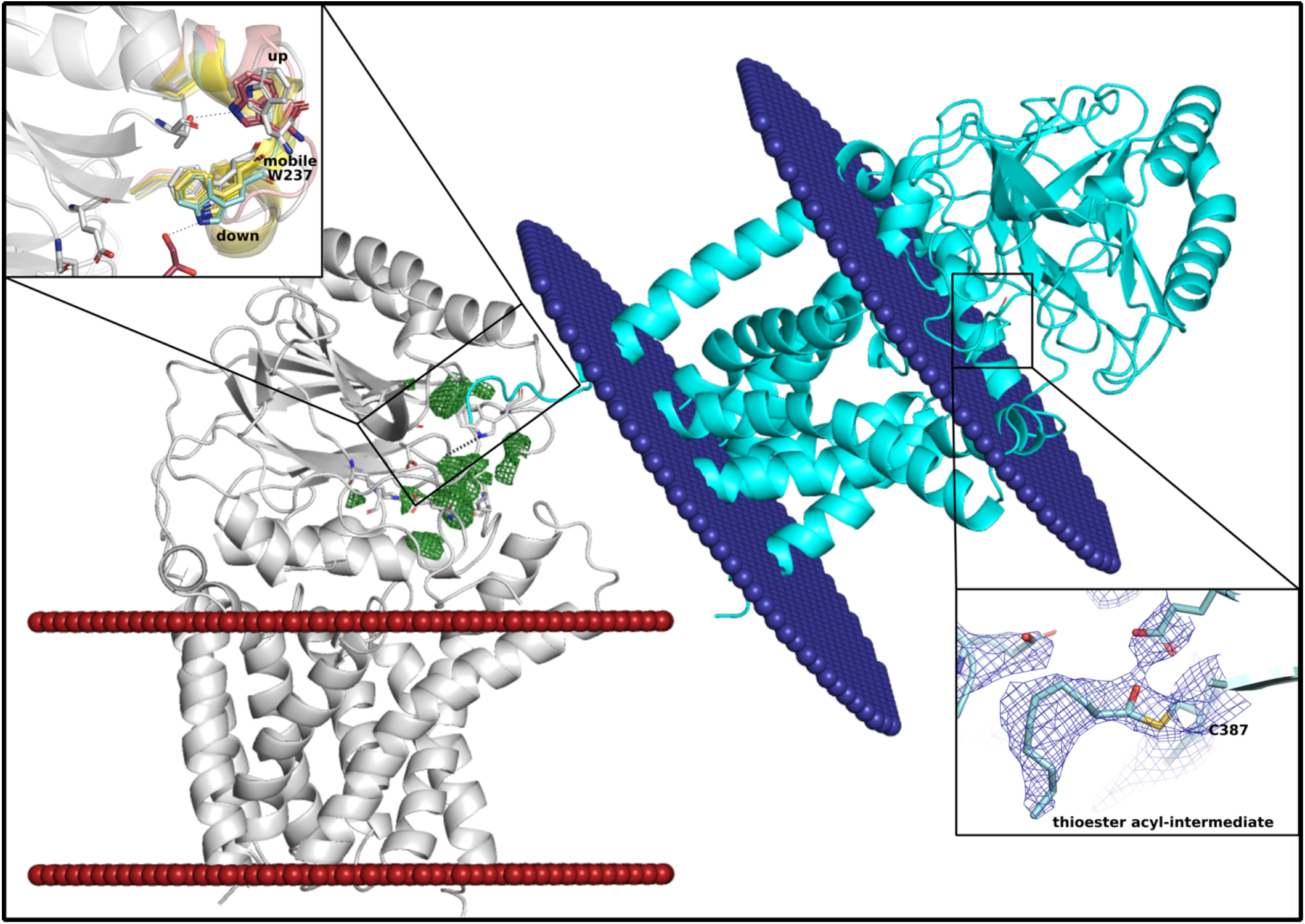

## INTRODUCTION

The post-translational modification of proteins by the covalent attachment of lipids is found across all domains of life. In bacteria, lipoproteins are important components of the cell envelope and are responsible for many essential cellular functions including nutrient uptake, secretion, cell wall integrity, and antibiotic production (Narita and Tokuda, 2017). In pathogenic bacteria lipoproteins are important virulence factors (Kovacs-Simon *et al.*, 2011). Pre-prolipoproteins are synthesized in the cytoplasm and translocated to the inner membrane via the Tat or Sec secretion pathways (Zückert, 2014). They contain an N-terminal signal peptide that contains a cysteine. Lipid attachment occurs at this cysteine via a sequential 3-step process (Figure 1A) controlled by three essential membrane bound enzymes, Prolipoprotein diacylglyceryl transferase (Lgt), Lipoprotein signal peptidase (LspA), and Apolipoprotein N-acyltransferase (Lnt) (Narita and Tokuda, 2017). The first step, carried out by Lgt, involves the transfer of an N-acyl diglyceride group to what will become the N-terminal cysteine. This is followed by the cleavage of the signal peptide by LspA to form apolipoproteins (Narita and Tokuda, 2017). The last step, unique to Gram negative bacteria, is the N-acylation of the terminal cysteine by Lnt to form the final mature lipoprotein (Hillmann *et al.*, 2011). The mature lipoprotein then remains at the inner membrane or is translocated to the outer membrane by the Lol pathway (Narita and Tokuda, 2011).

**Figure 1.**
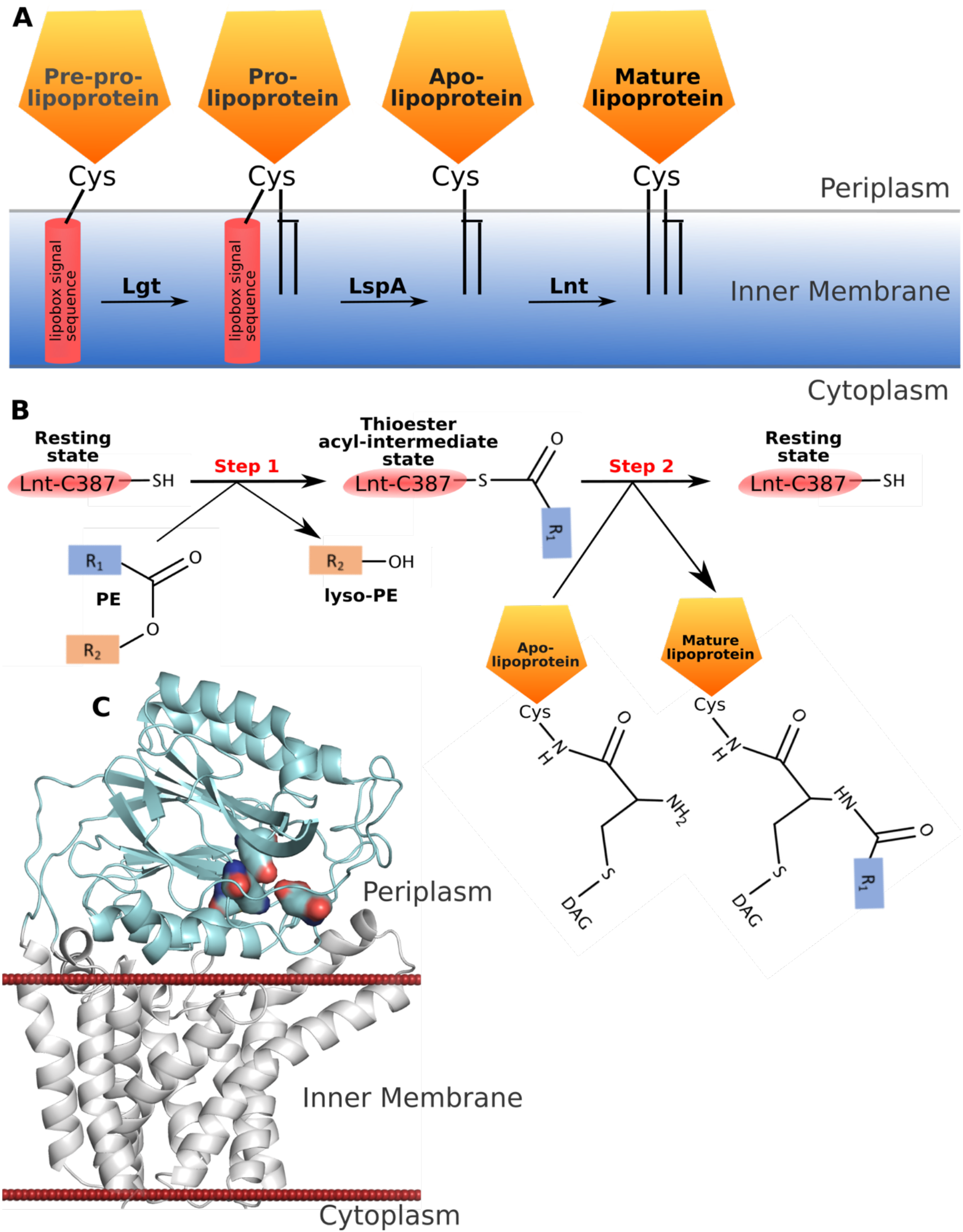
Mechanism and structure of Lnt. (A) Scheme of lipoprotein biosynthesis in Gram-negative bacteria. (B) The ping-pong reaction mechanism in Lnt. (C) Overall structure of Lnt consisting of 8 transmembrane helices (white) and periplasmic nitrilase domain (cyan). Surface representation of key residues, highlighting the position of the active site. The red spheres represent the position of the membrane as predicted by the PPM server (Lomize et al., 2012).

Based on sequence similarity Apolipoprotein N-acyltransferase (Lnt) belongs to the nitrilase superfamily. Nitrilases are multimeric proteins that contain a common Glu-Lys-Cys catalytic triad that hydrolyse carbon-nitrogen bonds (Bork and Koonon, 1994; Hung *et al.*, 2007; Weber *et al.*, 2013). In the case of Lnt, the nitrilase domain catalyzes the attachment of a phospholipid to the alpha-amino group of the N-terminal cysteine of the apolipoprotein creating the final mature lipoprotein. This attachment occurs via a proposed 2-step ping-pong mechanism (Wiktor *et al.*, 2017). The first step is the acyl transfer of the phospholipid substrate to create a thioester linkage on the active site cysteine. The second step is the transfer of the acyl chain from this cysteine to the N-terminal cysteine of the lipoprotein (Figure 1B).

All three enzymes involved in lipid attachment are essential for survival in bacteria making them attractive targets for new antimicrobial agents (Xia *et al.*, 2018). Recently both the structures of Lgt (Mao *et al.*, 2015) and LspA (Vogeley *et al.*, 2016) have been solved, and very recently a number of structures of Lnt became available (Noland *et al.*, 2017; Lu *et al.*, 2017; Wiktor *et al.*, 2017). Aside from the transmembrane domain, the structures of Lnt confirm that the overall fold of soluble nitrilases is conserved in Lnt. One key distinguishing feature of the Lnt nitrilase domain, however, is a long loop region that is longer and more flexible than is seen in typical soluble nitrilases that seems to extend parallel to the membrane in most cases (Gélis-Jeanvoine *et al.*, 2015, Cheng *et al.*, 2018). Overall, the reported structures of Lnt are all very similar and describe an enzyme essentially with or without its lipid substrate bound, with little or no information of how an incoming apolipoprotein peptide may interact with Lnt. Here we report the structure of the *E.coli* Lnt in its thioester acyl-intermediate state. We also report other substrate induced dynamics that may have important implications in active site access and catalysis.

## RESULTS AND DISCUSSION

### Overall Structure of Lnt

Data collection and refinement statistics are reported in Table 1. Briefly, we were able to crystallize two separate constructs of Lnt in two distinct crystallization conditions and crystal forms. Both forms have an overall architecture very similar to each other and to the recently reported structures with a transmembrane domain consisting of 8 helices and a soluble domain with a nitrilase fold (Figure 1C). Construct 1 (Lnt-C1) contains a fifteen-amino acid C-terminal tail that contains the His_8_-tag used for purification. It crystallized in space group P2_1_2_1_2_1_ using the vapor-diffusion method and contains two molecules in the asymmetric unit. Construct 2 (Lnt-C2) contains only two additional residues on its N-terminus that were left over after cleavage of the His_6_-tag used for purification. It crystallized in space group P6_4_22 using the lipidic cubic phase method and contains 1 molecule in the asymmetric unit. The structural differences between the protomers as well as the crystal packing properties of these two new crystal forms provide new insights into the mechanism and dynamics of this central enzyme.

### Structure of the thioester acyl-intermediate

As mentioned above Lnt-C1 crystallized with 2 molecules in the asymmetric unit. In one of those molecules (chain B) clear electron density can be seen extending off the active site C387 side chain that is not seen in the other molecule (Figure 2A). We interpret this as the thioester acyl-intermediate. This is entirely consistent with mass spectrometry analysis indicating that a large percentage of the protein is covalently bound to palmitate (Noland *et al.*, 2017) and previous findings that Lnt exists as a thioester acyl-enzyme intermediate (Buddelmeijer and Young, 2010). Despite seeing a significant percentage of covalently modified enzyme no electron density is seen in the structures presented by Noland *et al.*, 2017 or in the other solved structures of Lnt (Lu *et al.*, 2017; Wiktor *et al.*, 2017). Thus, this would be the first crystallographic evidence of the thioester acyl-enzyme intermediate. Also, since the protein was not crystallized in the lipidic cubic phase or supplemented with lipids during purification or the crystallization process it is likely that this modification is the natural intermediate state of the enzyme.

**Figure 2.**
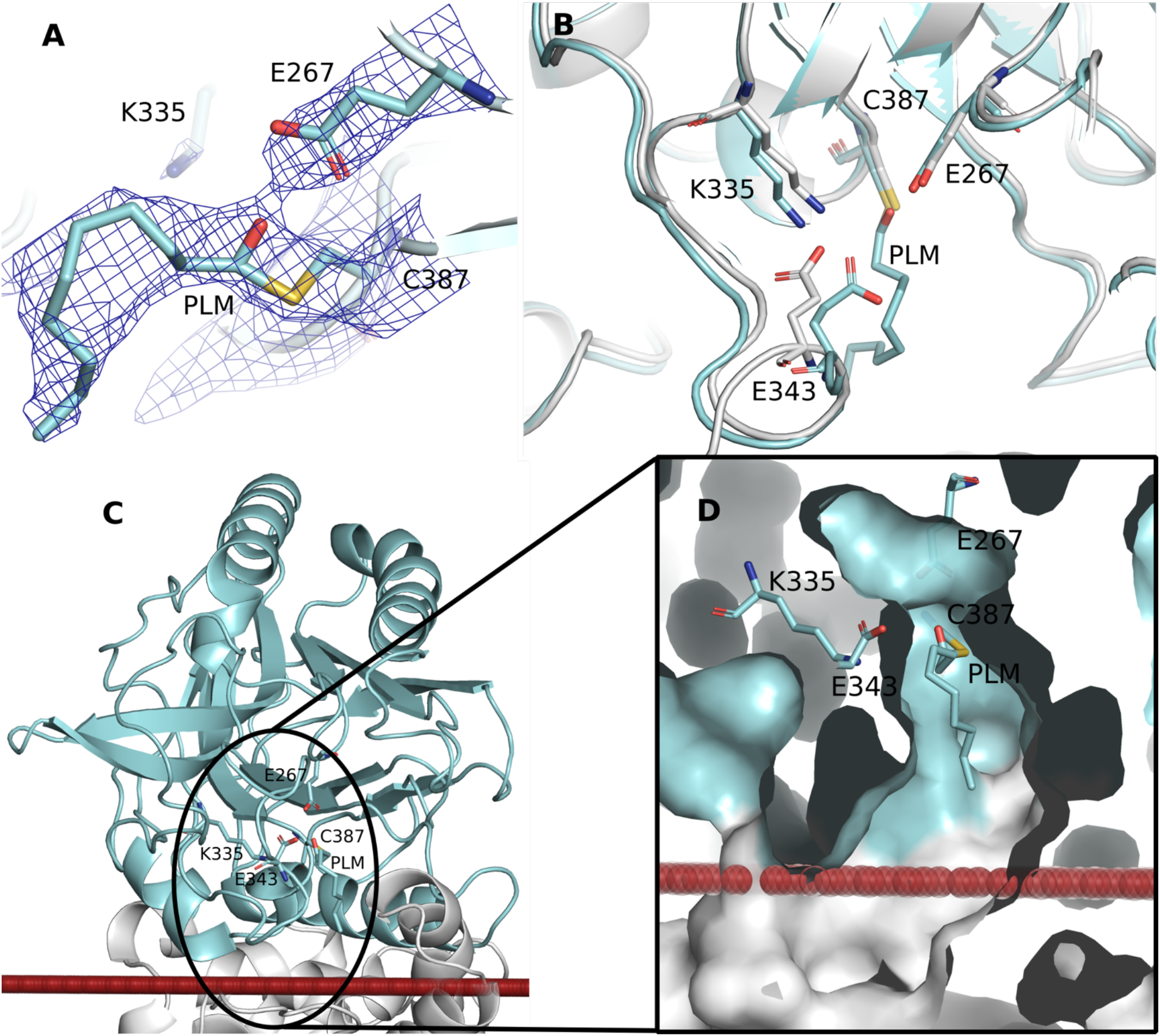
The thioester acyl-intermediate. (A) The active site of chain B and corresponding electron density modelled at 3σ showing the thioester acyl-intermediate. (B) Overlay of the active sites of chains A and B. Cartoon (C) and surface (D) representation of the large, open substrate entry portal leading to the active site in relation to the membrane interface (red spheres).

Remarkably, despite this modification, no large movements of side chains are required to accommodate this extra density compared with the other available models and the other (non-palmitated) molecule (chain A) in the asymmetric unit (Figure 2B). The catalytic cysteine, C387, of Lnt sits just above the membrane and at the back of a large open cavity (Figure 2A-C) that provides ample room for a palmitate molecule without the need for rearranging of the active site residues. There is, however, a slight movement towards the palmitate molecule of the side chain of E343 that could help stabilize this intermediate. This residue is also located at the hinge of a flexible loop structure in the active site vicinity.

As shown in Figure 2D, the acyl tail of the intermediate is positioned above the predicted membrane interface. Although the tail could not be confidently built into where the membrane region is expected to be, it is reasonable to assume that it extends into the membrane until the bottom of the binding pocket. It has also been suggested that Lnt exists at a slight angle with the binding pocket angling slightly downwards, towards the lipid interface (Lu *et al.*, 2017), thus the membrane interface could be higher than predicted by the PPM server. Regardless of the exact position of the membrane interface in relation to the binding pocket, the fact that the acyl tail can exist above the membrane interface can be easily explained by the high hydrophobicity of the binding groove (Noland *et al.*, 2017) that could easily accommodate hydrophobic molecules such as lipids.

### Dynamics of the 345-365 Arm

Together with the thioester acyl-intermediate, one of the striking features chain B of the Lnt-C1 structure is the position of the long 345-365 arm that is at an upward 60° angle to the membrane (Figure 3). In chain A by contrast, although residues 352-362 were not visible in the electron density, the beginning and end of the arm can be modelled extending parallel to the membrane (Figure 3B) similar to the other reported structures (pdb ids 5n6h, 5n6l, 5n6m, 5vrg, and 5vrh). When in this conformation, the active site is accessible to the surrounding environment, which could explain why the thioester intermediate is not seen in chain A or any of the other crystal structures despite mass spectral evidence (Noland *et al.*, 2017). However, this loop, when in the upward position could offer some protection to the active site and thioester intermediate from the outside environment.

**Figure 3.**
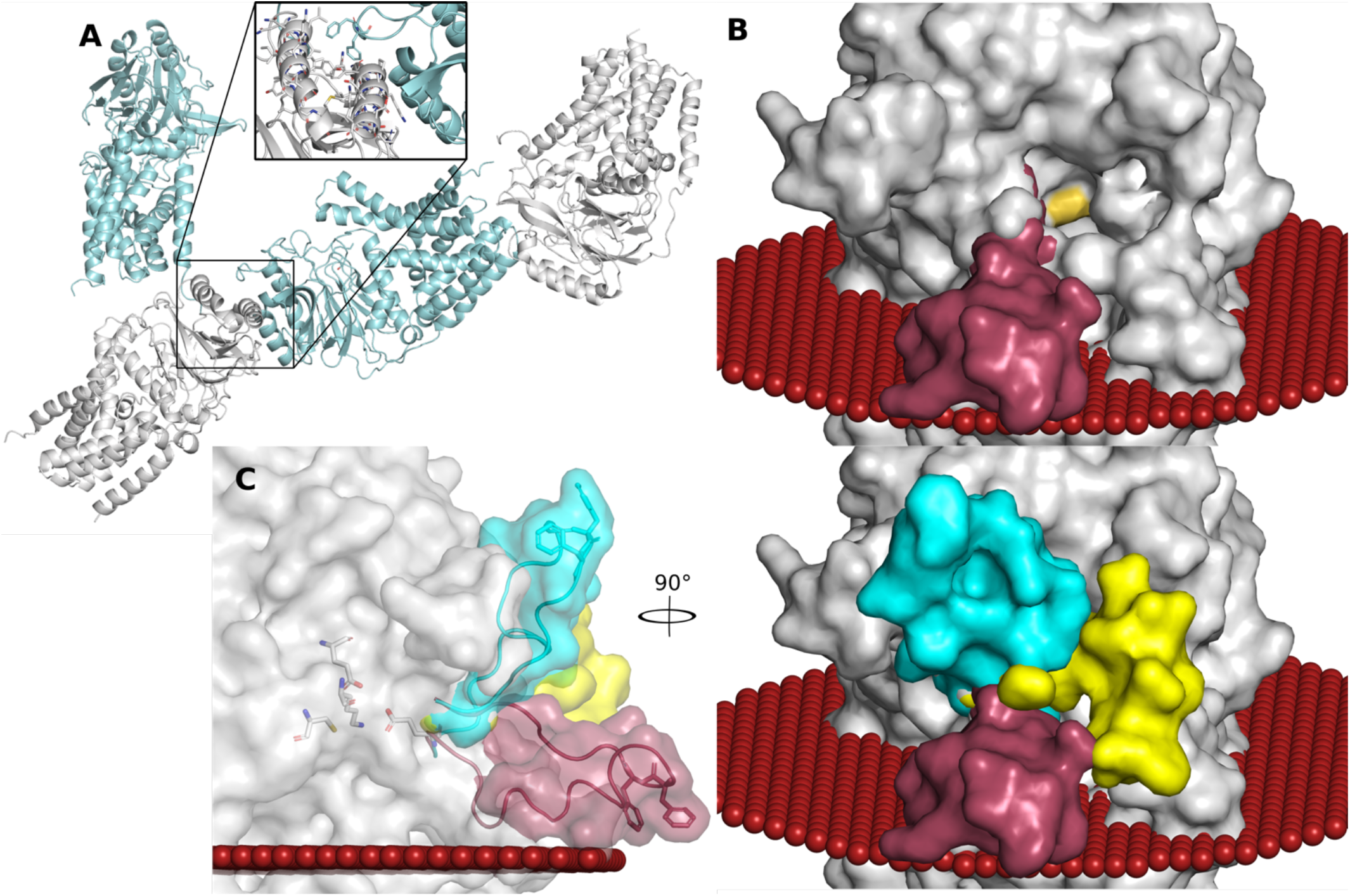
Dynamics of the flexible 345-365 arm. (A) P2_1_2_1_2_1_ crystal packing. Inset: Crystal contacts of F357 and F358 with an adjacent chain A molecule. (B) Surface representation of chain A (grey) and overlay of the flexible loop in an open conformation from pdb 5n6h (maroon) opening the substrate entry portal to the environment. Yellow patch is the location of the active site C387. (C) Surface representation of chain A overlaid with the various arm positions: open (maroon, pdb 5n6h), closed (cyan, chain B), middle (yellow, pdb 5xhq).

A second consequence of the mobile nature of this arm is the movement of the essential residue E343 that is at one of the joints allowing movement between the 2 conformations (Figure 2B), forming a salt bridge with the active site K335 when in the downward conformation, and pulling away from and breaking the salt bridge when in the upward position potentially stabilizing the thioester intermediate. This could provide long-range communication between the active site and the outer tip of the loop.

In total, 3 conformations of this arm have now been observed (Figure 3C) demonstrating its flexibility. In all cases where Lnt was crystallized in a more physiological environment using the LCP technique, the arm is positioned roughly parallel to the membrane interface. However, when crystallized using the vapor-diffusion and thus in the absence of a stabilizing lipid bilayer, (pdb id 5XHQ and chain B of Lnt-C1) this arm is noticeable higher than the predicted membrane interface. Molecular dynamics simulation (Lu et al, 2017) have suggested that this arm interacts strongly with the lipid bilayer consistent with the observation that all of the structures solved using the LCP method had this arm in a position that would be embedded in the monoolein membrane.

Interestingly, the only outlier to this trend is the protomer in chain A of Lnt-C1. Despite crystalizing using the vapor diffusion method its arm is positioned roughly parallel to the membrane similar to those crystalized using the LCP method. This might be explained by the arrangement of the protomers within the asymmetric unit (Figure 4C). The detergent belt of the chain B protomer within the asymmetric unit could be holding this arm down in this position.

The positioning of this arm is made possible by an adjacent chain A protomer that makes crystal contacts with F357 and F358 that pin the arm in this upward position. Despite F357 and F358’s location at the tip of this long arm, far removed from the active site, they have been previously shown to be essential for Lnt activity (Gélis-Jeanvoine *et al.*, 2015). Similarly, in the only other structure that a complete arm was able to be build (Wiktor *et al.*, 2017) F357 and F358 also form crystal contacts far removed from the active site. Thus, F357 and F358 could be used for directing and signaling to the active site through E343 about an incoming apolipoprotein molecule. Overall, the large degree of movement seen in this arm demonstrates its flexibility and ability to exist in and out of the membrane interface. These characteristics would be very beneficial in directing and stabilizing the various shapes and sizes of incoming substrate apolipoprotein molecules.

**Figure 4.**
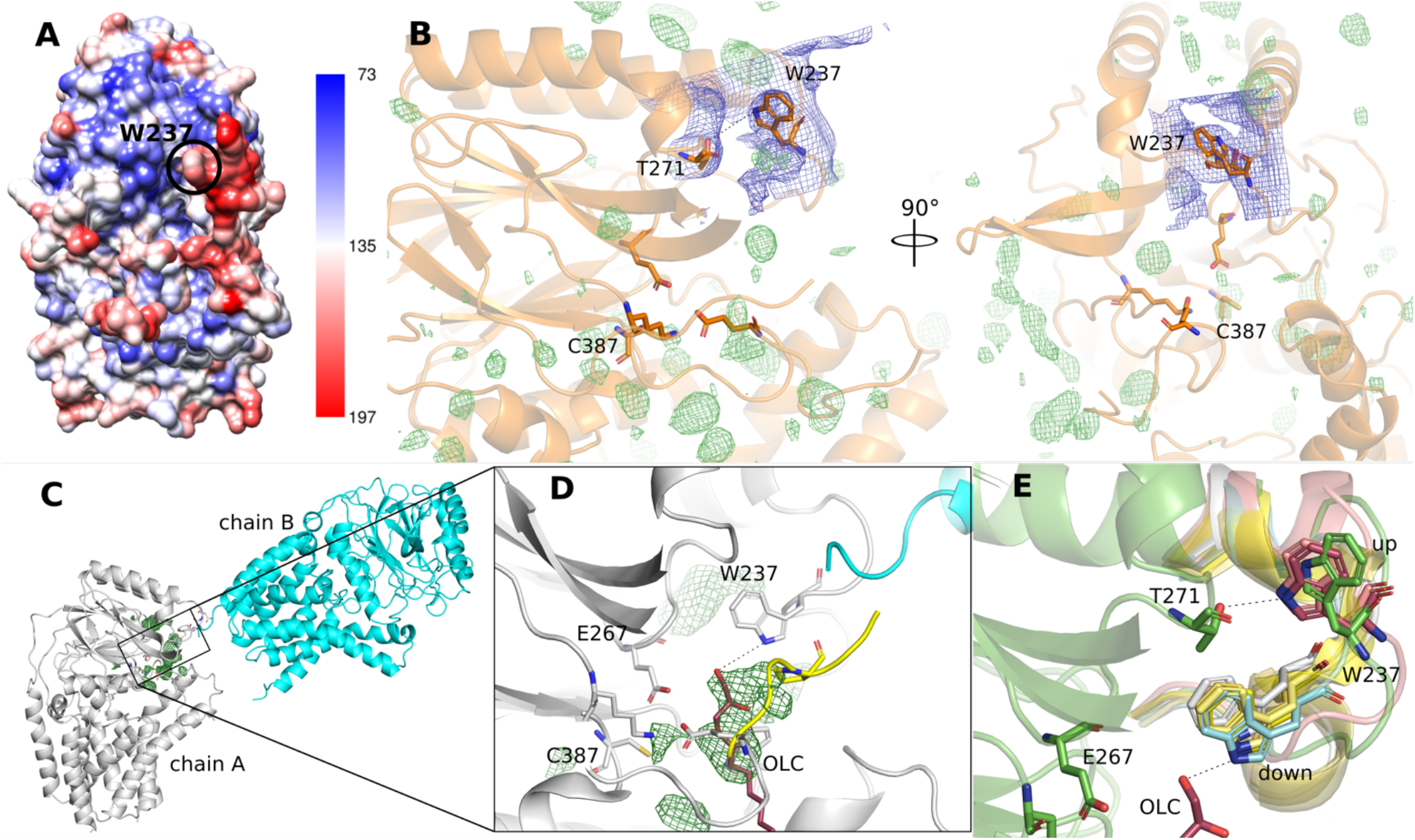
Substrate binding and structure of Lnt-C2. (A) Surface representation of Lnt-C2 colored according to B-factor. (B) In this crystal form W237 is in an upward position coordinated with T271. The substrate entry portal and active site are remarkably empty. Positive mF_o_-DF_c_ electron density modelled at 3σ (green). Electron density in the area of W237 modelled at 0.5σ (blue). (C) The asymmetric unit of Lnt-C1. A portion of the c-terminus from chain B (cyan) is positioned at the entry to the substrate portal of chain A (white). Green mesh represents unknown positive electron density modelled at 3σ at the entry to the substrate portal. (D) Zoom of boxed region in C. Overlay of OLC (red, pdb 5vrg) and a piece of the flexible 345-365 loop (yellow, pdb 5xhq) in the same position as unknown density. The essential W237 is in close coordination. (E) Position of W237 in the available crystal structures. In coordination with T271 when in the up position. In coordination with a lipid substrate when in the down position.

### Structure of Lnt in an apo-state

As mentioned above Lnt-C2 crystallized using the lipidic cubic phase method with 1 molecule in the asymmetric unit. Similar to Lnt-C1 the full 345-365 arm could not be built but appears to extend parallel to the membrane similar chain A of Lnt-C1 and other Lnt structures crystallized using the LCP method. The crystal structure of Lnt-C2 is remarkable in the fact that despite crystallizing in the lipidic cubic phase, it’s substrate binding portal is devoid of any bound substrates. In this crystal form, despite high B-factors in this area, W237 can be seen in the upward position coordinated to T271 (Figure 4A-B). Also, despite crystallizing in the presence of 400 mM Ammonium phosphate, no phosphate molecules could confidently be built into the model. In particular, Noland *et al* (2017) suggest a phosphate binding site at W237 based on binding of a chloride ion in that position, however we do not see density to support binding in this position. Moreover, the loop containing W237 has by far the highest B-factor of the entire protein (Figure 4A), if phosphate or other molecules are indeed bound there a stabilizing effect would be expected.

### Substrate mediated movement of W237

W237 has previously been shown to be essential for only the second step of the Lnt mechanism, i.e. the transfer of the acyl chain from Cys387 to the N-terminal cysteine of the lipoprotein (Gélis-Jeanvoine *et al.*, 2015).

As mentioned Lnt-C1 crystallized using the vapor diffusion technique with two molecules in the asymmetric unit. As shown in Figure 4C, the two molecules are arranged within the asymmetric unit in such a way that could give insights into one possible mode of apolipoprotein docking to Lnt. The C-terminus of chain B is in close proximity to the essential W237 at the entry of the substrate portal. Also, at the entry to the substrate portal, unexplained density is observed. Overlay of the pdb 5vrg places the head group of a molecule of monoolein into this density that would be coordinated to the nitrogen atom of W237. Similarly, overlay of the pdb 5xhq places a piece of the flexible 345-365 loop near this density that could also interact with W237 (Figure 4D).

W237 appears to be positioned at the entry of this large cavity leading to the active site triad. An overlay of all available crystal structures results in W237 in 2 alternate positions: an upward position interacting with T271 pointing away from the substrate portal, and a downward position pointing into the substrate portal allowing interaction with substrates (either protein or lipid) (Figure 4E). In the other crystal structures where W237 is in the upward position (pdb 5n6h and 5n6l) no substrates are seen in the substrate portal. In all cases where W237 is in the downward position, it is coordinated with a potential substrate (a lipid and/or peptide in the case of 5vrg, 5vrh, 5xhq, Lnt-C1). This suggests that substrate binding in the entry portal triggers movement to the downward position of W237, potentially triggering catalysis. It has also been suggested that F82 (Noland *et al.*, 2017), that is positioned opposite W237 also at the entrance to the portal acts as a potential gatekeeper. This cannot be ruled out, although unlike W237, an overlay of the available structures does not reveal a clear trend in the presence or absence of a substrate.

Together, the data presented here reveal a potential mode of apolipoprotein docking and points to an important role of substrate-induced protein dynamics for active site access and catalysis in apolipoprotein N-acyltransferase.

## Supporting information

