## Supplementary material for "Conformational changes in Apolipoprotein N-acyltransferase (Lnt)"

### STAR ★ METHODS

#### KEY RESOURCES TABLE

| REAGENT or RESOURCE | SOURCE | IDENTIFIER |
|---|---|---|
| Software, Algorithms, and Servers | | |
| XDS / XDSCONV | (Kabsch, 2010) | |
| PHENIX | (Adams et al., 2010) | |
| PHASER | (McCoy et al., 2007) | |
| COOT | (Emsley et al., 2010) | |
| MolProbility | (Chen et al., 2010) | http://molprobity.biochem.duke.edu/ |
| PyMol | Schrödinger LLC | http://pymol.org |
| UCSF Chimera | (Pettersen et al., 2004) | https://www.cgl.ucsf.edu/chimera/ |
| Protein Data Bank (PDB) | | http://pdb.org |
| PPM server | (Lomize et al., 2012) | http://opm.phar.umich.edu/server.php |
| Inkscape | | https://inkscape.org/ |
| Bacterial Strains and Vectors | | |
| *E. coli*: TOP10 | ThermoFisher | Cat. C404010 |
| Oligonucleotide primers | eurofins Genomics | |
| *E. coli*: C41 | Lucigen | Cat. 60442-1 |
| Vector for expression of C-terminal His-tag construct | (Sjöstrand et al., 2017) | |
| Vector for expression of N-terminal His-tag pET-28a | Novagen | Cat. 69864-3 |
| Chemicals and Recombinant Proteins | | |
| TEV protease | Sigma-Aldrich | Cat. T4455-1KU |
| n-nonyl- ß-D-maltoside (NM) detergent | Anatrace | Cat. N330 |
| n-dodecyl-ß-D-maltoside (DDM) detergent | Anatrace | Cat. D310 |
| Monoolein (9.9 MAG) | Molecular Dimensions | Cat. MD2-68 |
| Sodium Cacodylate | Hampton Research | Cat. HR2-239 |
| PEG 2000 MME | Molecular Dimensions | Cat. MD2-250-17 |
| PEG 200 | Molecular Dimensions | Cat. MD2-250-1 |
| Ammonium phosphate dibasic | Sigma-Aldrich | Cat. 379980-25G |
| Luria Broth (LB) growth media | Formedium | Cat. LBO0102 |
| MemMeso crystallization screen | Molecular Dimensions | Cat. MD1-86 |
| MemGold crystallization screen | Molecular Dimensions | Cat. MD1-74 |
| MemGold2 crystallization screen | Molecular Dimensions | Cat. MD1-74 |
| Other | | |
| Ni-NTA agarose | Qiagen | Cat. 30210 |
| Superdex 200 16/60 PG | GE Healthcare | Cat. 28-9893-35 |
| Emulsifex cell disrupter | Avestin | |
| LEX bioreactor | epiphyte3 | |
| Mosquito LCP dispensing robot | ttplabtech | |
| PDB Codes | | |
| *E. coli* Lnt | (Wiktor et al., 2017) | PDB: 5N6H, 5N6L |
| *P. aeruginosa* Lnt | (Wiktor et al., 2017) | PDB: 5N6M |
| *E. coli* Lnt | (Noland et al., 2017) | PDB: 5VRH, 5VRG |
| *E. coli* Lnt | (Lu et al., 2017) | PDB: 5XHQ |
| Deposited Data | | |
| Crystal structure of C-terminal His construct | This work | PDB: xxxx |
| Crystal structure of N-terminal His, TEV cleaved construct | This work | PDB: yyyy |

#### CONTACT FOR REAGENT AND RESOURCE SHARING

Further information and requests for resources and reagents should be directed to and will be fulfilled by Martin Högbom

#### METHOD DETAILS

##### Cloning, Expression and Protein Purification

The full length Lnt gene was amplified from *E.coli* K12 genomic DNA using standard PCR techniques with the primers 5' ACTCAGCTCGAGATGGCTTTTGCCTCATTAATTGAACGC 3' and 5' ATCGACGAATTCTTTACGTCGCTGACGCAGACTCATCAAC 5' and inserted into a modified pWaldo vector lacking the GFP reporter for a C-terminal $His_8$-tag construct (C1) (Sjöstrand *et al.*, 2017) using the restriction sites *XhoI* and *EcoRI*. The second construct (C2) was constructed using the primers 5' ACTCAGCTCCATATGGCTTTTGCCTCATTAATTG 3' and 5' ACTGACGAATTCTTATTTACGTCGCTGACGCAG 3' and inserted into the pET-28 vector (Novagen) containing a N-terminal TEV protease cleavable $His_6$-tag using the restriction sites *NdeI* and *EcoRI*.

For large-scale protein purifications, a plasmid containing Lnt was transformed into chemically competent *E.coli* C41. A pre-culture from a single colony was grown overnight at 37 °C in LB broth containing 50 µg/mL kanamycin, which was then used to inoculate 1.5 L LB broths in a LEX bioreactor (epiphyte3). Cells were grown at 37 °C until an $OD_{600}$ = 1.0 prior to the induction of protein expression with 0.4 mM IPTG at ambient temperature for 4 hours. Cells were then harvested *via* centrifugation at 6000 g for 15 minutes and frozen at -20 °C. Frozen cells were thawed and suspended in lysis buffer (50 mM TRIS pH 8.0), and lysed by passage of at least 3 times through an emulsifex cell disrupter (Avestin). The lysate was centrifuged at 16000 g for 40 minutes to pellet cell debris. Membranes were then isolated by ultra-centrifugation of the cleared lysate at 40000 g for 40 minutes; washed one time with a washing buffer (500 mM NaCL, 50 mM TRIS pH 8.0); ultra-centrifuged again, and finally suspended in buffer containing 300 mM sucrose, 20 mM TRIS, pH 8.0, and frozen at -80 °C.

Purified membranes were diluted 20 times with buffer A (50 mM TRIS pH 8.0, 400 mM NaCl, 5 % glycerol, 10 mM imidazole, 5 mM β-mercaptoethanol) and solubilized with 1 % n-Dodecyl-β-D-Maltopyranoside (DDM) (Anatrace) at 4 °C for 90 minutes on a mixer, followed by centrifugation at 40000 g for 45 minutes to remove unsolubilized membranes. The resulting supernatant was incubated at 4 °C on a mixer with Ni-NTA resin (Qiagen) equilibrated with buffer A supplemented with 0.05 % DDM for 90 minutes. The resin was washed with 100 mL buffer B (50 mM TRIS pH 8.0, 200 mM NaCl, 5 % glycerol, 25 mM imidazole, 5 mM β-mercaptoethanol, 0.04 % DDM), followed by a second 100 mL of buffer C (50 mM TRIS pH 8.0, 200 mM NaCl, 5 % glycerol, 50 mM imidazole, 5 mM β-mercaptoethanol, 0.04 % DDM). The C-terminal construct was eluted from the Ni-NTA resin with buffer D (50 mM TRIS pH 8.0, 150 mM NaCl, 5 % glycerol, 250 mM imidazole, 5 mM β-mercaptoethanol, 0.4 % n-Nonyl-β-D-Maltopyranoside (NM). The eluted protein was concentrated with a 100 MWCO ultrafiltration spin column (Vivaspin) to less than 5 mL and further purified by injection into a HiLoad 16/60 Superdex 200 column (GE Healthcare) equilibrated with buffer E (25 mM TRIS pH 8.0, 100 mM NaCl, 5 % glycerol, 0.5 mM TCEP, 0.35 % NM). The purified protein eluted as a single peak at approximately 73 mL volume. The appropriate fractions were pooled and concentrated with a 100 MWCO to 15 mg/mL. The final purified protein was snap frozen in liquid nitrogen, and stored at -80 °C if not used immediately.

The N-terminal TEV-cleaved construct (C2) was purified as described above except the protein was eluted from the Ni-NTA resin with buffer F (50 mM TRIS pH 8.0, 150 mM NaCl, 5 % glycerol, 250 mM imidazole, 5 mM β-mercaptoethanol, 0.03 % DDM). The eluted protein was concentrated with a 100 MWCO ultrafiltration spin column (Vivaspin) to less than 5 mL, diluted with buffer G (25 mM TRIS pH 8.0, 100 mM NaCl, 5 % glycerol, 0.5 mM TCEP, 0.02 % DDM) tenfold and incubated at 4 °C with TEV protease containing a $His_6$-tag in a 1:10 ratio overnight on a mixer. The next morning, the mixture was then incubated at 4 °C with Ni-NTA resin equilibrated with buffer G for 2 hours on a mixer. The Ni-NTA resin was collected with a gravity flow column and the flow-through was concentrated to less than 5 mL and further purified by injection into the HiLoad 16/60 Superdex 200 column (GE Healthcare) equilibrated with buffer G. The TEV-cleaved purified protein eluted as a single peak at approximately 70 mL volume. The appropriate fractions were pooled and the final purified protein was concentrated with a 100 MWCO filter to 15 mg/mL, snap frozen in liquid nitrogen, and stored at -80 °C for subsequent use in crystallization assays.

**Crystallization**

For crystallization trials with the C-terminal protein at 15 mg/mL was used to screen for initial crystal hits using the commercially available screens Memgold and Memgold2 (Molecular Dimensions). Crystallization trials were set up using a Mosquito liquid dispenser robot (ttplabtech) into 96 well, 2 drop, vapor diffusion sitting drop plates with 50 µL reservoir solution and 1:1 and 2:1 protein:precipitate ratios. The final optimized condition was 25 % PEG2000MME, 100 mM Na cacodylate pH 6.4, 40 mM $MgCl_2$ grown at 18 °C, with a drop ratio of 1:1.75 protein:precipitate. Crystals grew to about 0.3 x 0.2 x 0.2 mm in about 2 weeks and were loop-harvested and snap-cooled in liquid nitrogen without added cryo-protectant.

For crystallization trials with the N-terminal, TEV cleaved protein, Lnt was reconstituted into the lipidic cubic phase using standard techniques (Caffrey and Cherezov, 2009). The protein solution at 15 mg/mL was homogenized with monoolein (9.9 MAG) in a coupled syringe mixing device using two volumes of protein solution and three volumes of lipid. Initial crystallization trials were set up by transferring 50 nL of the protein-laden mesophase onto a silanized 96-well glass sandwich plate followed by 800 nL of precipitant solution using a LCP-Mosquito liquid dispenser robot (ttplabtech) and stored at 18 °C for crystal growth. Optimized crystals were obtained with 36 % PEG200, 400 mM Ammonium phosphate dibasic, 100 mM HEPES pH 7.2. High diffracting crystals were only able to be obtained by storing the glass plate at 10 °C for 3 weeks, followed by transfer to 4 °C for an additional 3 weeks. Crystals were loop harvested and snap frozen in liquid nitrogen without the addition of cryo-protectant after 6 weeks of growth.

**Data collection, Processing, and Structure Determination**

During the optimization process X-ray diffraction experiments were carried out at various synchrotron light sources across Europe, namely the ESRF, DIAMOND, BESSY II, and the SLS. The final high resolution datasets were collected at the SLS, beamline X06SA-PX. Data was processed with XDS (Kabsch, 2010). The structures were solved by molecular replacement with the pdb model 5N6L as a search model searching for 2 molecules in the asymmetric unit for the C-terminal construct and 1 molecule for the N-terminal TEV cleaved construct using the PHENIX suite of programs (Adams *et al*., 2010) with final model building in Coot (Emsley and Cowtan, 2004). Data collection parameters and refinement statistics are reported in Table 1. Figures were generated using PyMOL (Delano, 2010) or UCSF Chimera (Pettersen *et al*., 2004).

#### DATA AND SOFTWARE AVAILIBILITY

##### Accession Numbers

The atomic coordinates and structure factors for the structures described in this study have been deposited to the RCSB PDB (www.rcsb.org) with the PDB ID codes: xxxx and yyyy.

#### ADDITIONAL RESOURCES

N/A

**Table 1. Data Collection and Refinement Statistics**

| | Lnt-C1<br>PDB: xxxx | Lnt-C2<br>PDB: yyyy |
|---|---|---|
| Data Collection | | |
| X-ray source | SLS, X06SA-PX | SLS, X06SA-PX |
| Wavelength (Å) | 1.00 | 1.00 |
| Space group | $P2_12_12_1$ | $P6_422$ |
| Cell Dimensions | | |
| a, b, c (Å) | 72.28, 137.287, 221.236 | 80.725, 80.725, 442.575 |
| α, β, γ (°) | 90.0, 90.0, 90.0 | 90.0, 90.0, 120 |
| Resolution (Å) | 48.56-3.3 (3.4-3.3) | 46.89-3.01 (3.2-3.01) |
| Completeness | 99.8 (100) | 99.7 (100) |
| Redundancy | 7.3 (7.7) | 20.9 (19.2) |
| $I/\sigma(I)$ | 6.38 (0.63) | 13.97 (0.59) |
| $R_{meas}$ | 0.0790 (1.22) | 0.194 (4.93) |
| $CC_{1/2}$ | 0.998 (0.182) | 0.999 (0.232) |
| Refinement | | |
| Resolution (Å) | 48.56-3.3 | 46.89-3.1 |
| No. of unique reflections | 33906 (3299) | 29269 (2907) |
| $R_{work}/R_{free}$ | 0.27/0.31 | 0.26/0.30 |
| No. of atoms | 8032 | 3920 |
| Protein | 7918 | 3877 |
| Ligands | 114 | 43 |
| B-factors (Å$^2$) | | |
| Protein | 134 | 120 |
| Ligands | 180 | 129 |
| RMSD | | |
| Bond length (Å) | 0.002 | 0.004 |
| Bond angles (°) | 0.56 | 0.96 |
| Ramachandran plot (%) | | |
| Favored | 93.18 | 94.26 |
| Allowed | 6.72 | 5.74 |
| Outliers | 0.10 | 0.00 |

Values in parentheses are for the highest-resolution shell. RMSD, root-mean-square deviation
